# Identification of a Small Molecule Inhibitor of the Aminoglycoside 6’-*N*-Acetyltransferase Type Ib [AAC(6’)-Ib] Using Mixture-Based Combinatorial Libraries

**DOI:** 10.1101/198176

**Authors:** Tung Tran, Kevin Chiem, Saumya Jani, Brock A. Arivett, David Lin, Rupali Lad, Verónica Jimenez, Mary B. Farone, Ginamarie Debevec, Radleigh Santos, Marc Giulianotti, Clemencia Pinilla, Marcelo E. Tolmasky

## Abstract

The aminoglycoside 6′-*N*-acetyltransferase type Ib [AAC(6’)-Ib] is the most widely distributed enzyme among AAC(6’)-I-producing Gram-negative pathogens and confers resistance to clinically relevant aminoglycosides including amikacin. This enzyme is therefore ideal to target with enzymatic inhibitors that could overcome resistance to aminoglycosides. The search for inhibitors was carried out using mixture-based combinatorial libraries, the scaffold ranking approach, and the positional scanning strategy. A library with high inhibitory activity had pyrrolidine pentamine scaffold and was selected for further analysis. This library contained 738,192 compounds with functionalities derived from 26 different amino acids (R1, R2 and R3) and 42 different carboxylic acids (R4) in four R group functionalities. The most active compounds all contained S-phenyl (R1 and R3) and S-hydromethyl (R2) functionalities at three locations and differed at the R4 position. The compound containing 3-phenylbutyl at R4 (compound **206**) was a robust enzymatic inhibitor in vitro, in combination with amikacin potentiated the inhibition of growth of three resistant bacteria in culture, and improved survival when used as treatment of *Galleria mellonella* infected with *aac(6’)-Ib*-harboring *Klebsiella pneumoniae* and *Acinetobacter baumannii* strains.

## 1. Introduction

Drug resistance is a major clinical and public health problem. Several nosocomial and community acquired pathogens have become resistant to many different antibiotics, seriously complicating treatment and in some cases becoming virtually untreatable [1]. While there are numerous antimicrobials in development to treat drug resistant bacteria, the rate of introduction of new antibiotic classes in the clinics is being outpaced by the growth in multidrug resistant strains [1]. It is therefore desirable to find strategies to prolong the effectiveness of existing antibiotics. Aminoglycosides are mainly used to treat infections caused by Gram-negatives as well as Gram-positives in various combination therapies [2]. Additionally, these antibiotics are used as treatments of plague, tularemia, brucellosis, endocarditis and other infections caused by streptococci and enterococci, as well as *Mycobacterium tuberculosis* infections [2]. Aminoglycosides, in particular amikacin, are also administered to neonates as soon as Gram-negative infections are suspected because of the high morbidity and mortality that these infections have on this population [3]. Unfortunately, as it is the case with other kinds of antibiotics, aminoglycosides are losing their power due to the rise in the number of resistant strains [4, 5].

Resistance to aminoglycosides can occur through several mechanisms that can coexist simultaneously in the same cell [4, 5]. However, enzymatic inactivation of the antibiotic molecule is the most prevalent in the clinical setting. One of the most relevant aminoglycoside modifying enzymes is the aminoglycoside 6’-*N*-acetyltransferase type Ib [AAC(6’)-Ib] [2, 5, 6]. This enzyme, present in a vast majority of AAC(6′)-I-producing Gram-negative clinical isolates [2, 5, 6], confers resistance to amikacin and other important aminoglycosides. Of high relevance is its ability to inactivate amikacin, a semisynthetic derivative of kanamycin A that is one of the most refractory to the action of aminoglycoside modifying enzymes and has been instrumental in the treatment of severe multiresistant infections including life-threatening outbreaks in neonatal wards [3, 7, 8].

Although considerable efforts are dedicated to designing new aminoglycosides that can overcome existing mechanisms of resistance [4, 9], other alternatives to counter the action of aminoglycoside modifying enzymes are being researched. They include the utilization of inhibitors of gene expression such as antisense oligonucleotide analogs [10-12], inhibitors of the enzymatic activity like those developed to overcome β-lactamase-mediated resistance to β-lactams [5, 13-18], or metal ion inhibitors of the acetylation reaction, which most probably act by protective chelation of the substrate antibiotic [19-22]. However, in spite of the efforts directed at designing inhibitors of aminoglycoside modifying enzymes or their expression, none are in use in the clinics yet [reviewed in 4, 5]. Recognizing the clinical relevance of AAC(6’)-Ib, robust inhibitors of this enzyme were searched screening mixture-based combinatorial libraries using the scaffold ranking approach and the positional scanning strategy [23, 24]. This paper describes the identification of an inhibitor of AAC(6’)-Ib from a pyrrolidine pentamine scaffold library.

## 2. Materials and Methods

### 2.1 Bacterial strains

*A. baumannii* A155 and *K. pneumoniae* JHCK1 are multidrug resistant clinical strains that naturally harbor *aac(6’)-Ib* [7, 10]. *E. coli* TOP10(pNW1) is a laboratory strain obtained by transformation of *E. coli* TOP10 with pNW1, an F′ derivative including *aac(6’)-Ib* [25]. All four strains were used to test the ability of selected compounds to reduce the levels of resistance to amikacin. *E. coli* TOP10 harboring a recombinant clone where *aac(6’)-Ib* was placed under the control of the BAD promoter in the cloning vehicle pBAD102 to obtain a His-Patch-containing thioredoxin fused protein was used to purify the enzyme for enzymatic assays [15].

### 2.2 General methods

Purification of the enzyme was carried out as previously described [15]. Enzymatic activity was determined monitoring the increase in OD_412_ after the reaction of 5,5′-dithiobis(2-nitrobenzoic acid) (DTNB) with the CoA-SH released when acetyl CoA is used as donor for acetylation of the substrate aminoglycoside [16]. Libraries were dissolved in dimethylformamide (DMF) and the final concentration of DMF in the reaction mixtures used scaffold ranking was 9%. A control reaction was carried out in the presence of 9% DMF. Individual compounds were dissolved in dimethyl sulfoxide (DMSO) and the final concentration of DMSO in the reaction mixtures was 18%. Degree of acetylation was assessed monitoring OD_412_ in a BioTek Synergy 2 plate reader. Initial velocities (Vi) were calculated using the Gen 5 software, version 2.01.13. Apparent inhibition through protein aggregation was discarded by carrying out the reaction in the presence of Triton X-100 (0.1%) [15]. Results are averages of three separate experiments. Mode of inhibition was determined using Lineweaver–Burk plots and the Michaelis–Menten equation variations for the different types of inhibition using GraphPad Prism 6 software as before [14]. Datasets were generated performing a series of reactions in the presence of a range of inhibitor concentrations with one substrate at a constant excess concentration and the other at different concentrations. Inhibition of growth in the presence of amikacin and the testing compounds were carried out in Mueller-Hinton broth containing the indicated additions in a microplate reader (BioTek Synergy 5) as described before [20].

### 2.3 Libraries, synthesis, and purification of small molecule compounds

All mixture-based libraries screened were synthesized at Torrey Pines Institute for Molecular Studies using solid-phase chemistry approaches, simultaneous multiple syntheses, and “libraries from libraries” approaches, as previously described [24, 26-28]. The positional scanning library TPI1343 and all individual compounds reported here were synthesized using a previously described methodology [29]. Scheme 1 (Fig. S1, supplemental material) shows the general synthesis approach; a polyamide scaffold (Scheme 1; 2, Fig. S1, supplemental material) was synthesized on the solid support using standard Boc chemistry, the amide residues were then reduced with borane (Scheme 1;3, Fig. S1, supplemental material), and the compounds were then removed from the solid support (Scheme 1; 4, Fig. S1, supplemental material). The most active individual compounds (Table 1) were synthesized using the conditions described in Scheme 1 with each R position defined.

**Table 1.**
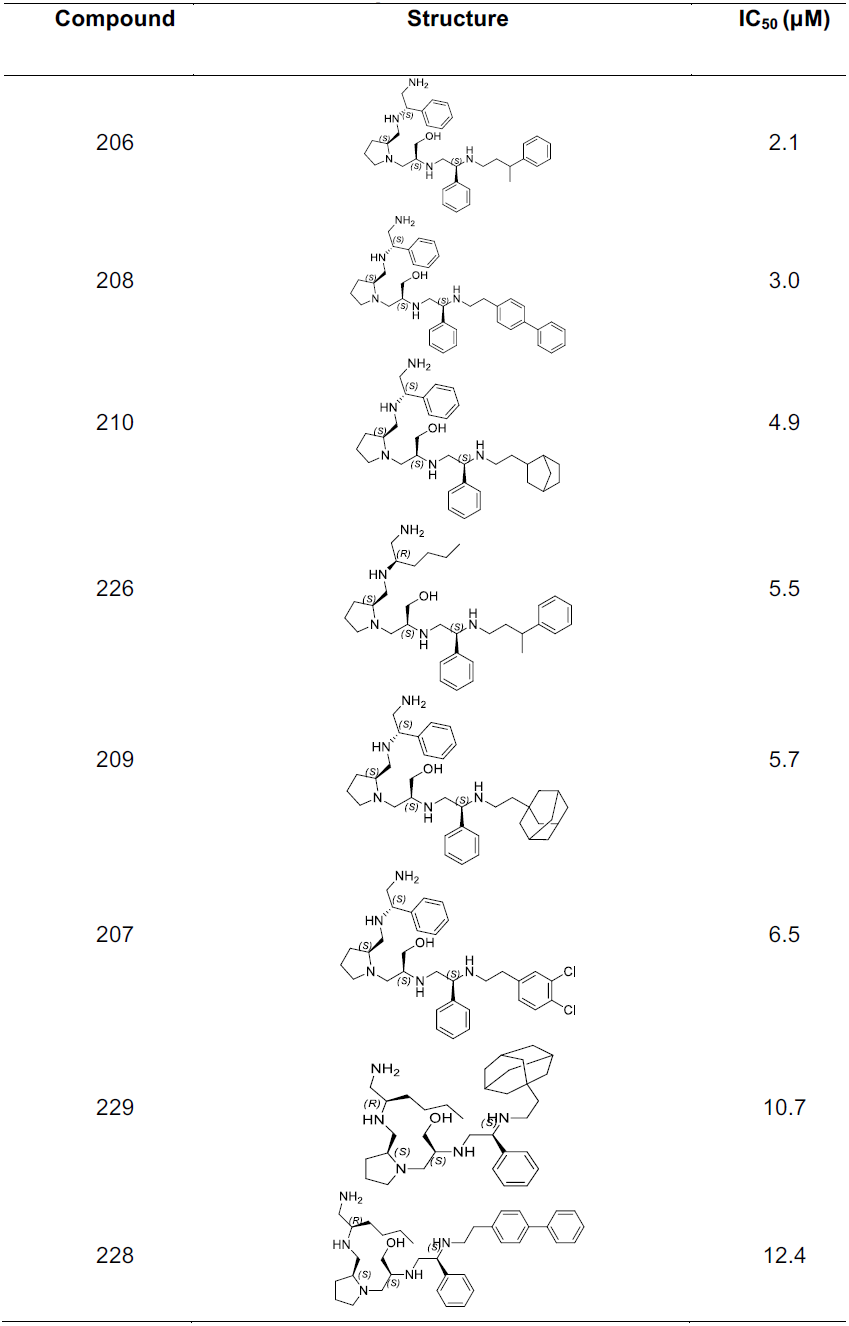
Most active compounds Compound Structure IC_50_ (µM)

| Compound | Structure | IC<sub>50</sub> (μM) |
| --- | --- | --- |
| 206 |  | 2.1 |
| 208 |  | 3.0 |
| 210 |  | 4.9 |
| 226 |  | 5.5 |
| 209 |  | 5.7 |
| 207 |  | 6.5 |
| 229 |  | 10.7 |
| 228 |  | 12.4 |

### 2.4 High-performance liquid chromatography (HPLC) purification

All purifications were performed on a Shimadzu Prominence preparative HPLC system, consisting of LC-8A binary solvent pumps, an SCL-10A system controller, a SIL-10AP auto sampler, and an FRC-10A fraction collector. A Shimadzu SPD-20A UV detector set to 214 nm was used for detection. Chromatographic separations were obtained using a Phenomenex Luna C18 preparative column (5 µm, 150 mm × 21.5 mm i.d.) with a Phenomenex C18 column guard (5 µm, 15 mm × 21.2 mm i.d.). Prominence prep software was used to set all detection and collection parameters. The mobile phases for HPLC purification were HPLC grade obtained from Sigma-Aldrich and Fisher Scientific. The mobile phase consisted of a mixture of acetonitrile/ water (both with 0.1% trifluoroacetic acid). The initial setting for separation was 2% acetonitrile, which was held for 2 min, then the gradient was linearly increased to 20% acetonitrile over 4 min. The gradient was then linearly increased to 55% acetonitrile over 36 min. The HPLC system was set to automatically flush and re-equilibrate the column after each run for a total of four column volumes. The total flow rate was set to 12 ml/min, and the total injection volume was set to 3900 µl. The fractions corresponding to the desired product were then combined and lyophilized.

### 2.5 Liquid chromatography–mass spectrometry (LCMS) analysis of purified material

Purity and identity of compounds was verified using a Shimadzu 2010 LCMS system, consisting of a LC-20AD binary solvent pumps, a DGU-20A degasser unit, a CTO-20A column oven, and a SIL-20A HT auto sampler. A Shimadzu SPD-M20A diode array detector scanned the spectrum range of 190-600 nm during the analysis. Chromatographic separations were obtained using a Phenomenex Luna C18 analytical column (5 µm, 50 mm × 4.6 mm i.d.) with a Phenomenex C18 column guard (5 µm, 4 × 3.0 mm i.d.). All equipment was controlled and integrated by Shimadzu LCMS solutions software version 3. Mobile phase A for LCMS analysis was LCMS grade water, and mobile phase B was LCMS grade acetonitrile obtained from Sigma-Aldrich and Fisher Scientific (both with 0.1% formic acid for a pH of 2.7). The initial setting for analysis was 5% acetonitrile (v/v), and then linearly increased to 95% acetonitrile over 6 min. The gradient was then held at 95% acetonitrile for 2 min before being linearly decreased to 5% over 0.1 min and held until stop for an additional 1.9 min. The total run time was equal to 12 min, and the total flow rate was 0.5 ml/min. The column oven and flow cell temperature for the diode array detector was 30°C. The auto sampler temperature was held at 15°C, and a 5 µl aliquot was injected for analysis.

### 2.6 ^1^H nuclear magnetic resonance (NMR) of purified compounds

^1^H NMR spectra were obtained utilizing the Bruker 400 Ascend (400 MHz). NMR chemical shifts were reported in δ (ppm) using the δ7.26 signal of CDCl_3_ or the δ2.50 signal of DMSO-*d*_*6*_ (^1^H NMR) as the internal standard (NMR Data, supplemental material).

### 2.7 Checkerboard assays and statistical analysis

Checkerboard assays were performed using variable titration of a given compound (in general, from zero to 24 µM in five doses) and amikacin (in general, six doses with a maximal dose of 16 µg/ml for *E. coli* TOP10(pNW1) and 64 µg/ml for the other strains). Statistical analysis was first done using the standard fractional inhibitory concentration (FIC) methodology [30]. However, due to the presence of compound activity a mixture modeling approach [31] that better quantifies exact levels of synergistic potentiation was developed and applied. In particular, assuming amikacin and the given compounds have independent mechanisms of action, one can model the percent activity of the mixture of the two substances as:

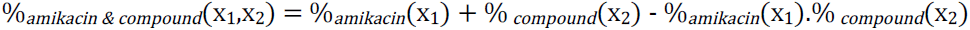

Here, x_1_ and x_2_ are the concentrations of amikacin and compound tested, respectively. This equation can be rearranged to model the effective percent activity of the antibiotic alone at a given concentration, after accounting for compound

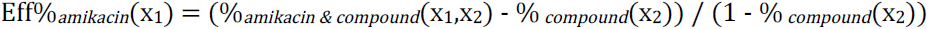

Thus, the model-adjusted checkerboards show the antibiotic activity post-potentiation, which permits determination of the true change in level of resistance.

### 2.8 Infection assays

Bacterial cells were collected by centrifugation and resuspended in phosphate-buffered saline (PBS) (7.2 pH) with the additions indicated in each assay. The number of cells inoculated was estimated based on the OD_600_ and the approximate number of bacterial cells inoculated was 5 x 10^5^ (±0.5 log) in a volume of 5 µl. The injections were performed using a syringe pump (New Era Pump Systems, Inc., Wantagh, NY) with a 26-gauge by half-inch needle into the hemocoel at the last left proleg after swabbing the zone with ethanol immediately prior to injection. Ten healthy randomly selected final-instar *G. mellonella* larvae (Grubco, Fairfield OH), weighing 250 mg to 350 mg, were used for each group (*n* = 30) and the experiments were carried out in triplicate. If more than two deaths occurred in either of the control groups, i.e., the PBS-injected group or the non-injected group, the entire trial was omitted. Larvae were incubated at 37°C for 120 h while in the dark while survival was recorded at 24-h intervals, removing dead larvae at reported intervals. The Kaplan-Meier method was used to plot resulting survival curves. A *P* value of 0.05 was considered statistically significant for the log-rank test of survival curves (Prism 6.0, GraphPad Software Inc., La Jolla, CA USA).

### 2.9 Cytotoxicity assays

Levels of cytotoxicity of compound **206** were determined using a commercial kit (LIVE/DEAD Viability/Cytotoxicity Kit for mammalian cells, Molecular Probes). HEK 293 cells were maintained at 37°C and 5% CO_2_ in high glucose DMEM media, supplemented with 10% fetal bovine serum. Cells were plated at a density of 1000 cells per well in flat bottom 96-well black microtiter plates and cultured overnight under standard conditions. Compound **206** was added to the cells at increasing concentrations and incubated for 24 hours. Equivalent concentrations of DMSO were used as controls. The cells were then washed with sterile D-PBS and incubated with the LIVE/DEAD reagent (2 µM ethidium homodimer 1 and 1 µM calcein-AM) for 30 min at 37°C. Fluorescence of the dyes was measured in a microtiter plate reader at 645 (dead cells) and 530 (live cells) nm respectively. The percentage of dead cells was calculated relative to the cells treated with DMSO. Cells incubated with 0.1% Triton X-100 for 10 min was used as a control for maximum toxicity. Experiments were conducted in triplicate. The results expressed as Mean ± SD of 3 independent experiments.

## 3. Results

The “scaffold ranking” approach was applied to initiate the identification of compounds that inhibit the AAC(6’)-Ib enzyme using a small molecule mixture based library in positional scanning format [24, 32]. The ranking was carried out testing 37 samples that contain a mixture of all the compounds for a given scaffold (Table S1, supplemental material). As described previously, these scaffold-ranking samples can be prepared as synthetic mixtures or by pooling together aliquots of each sample in a given positional scanning library [23, 29, 33].

The strategy used for identifying individual compounds from the combinatorial libraries is outlined in Fig. 1. The screening of the scaffold ranking samples for their inhibitory activity of AAC(6’)-Ib was carried out at 25 µg/ml. Fig. 2 shows the % inhibition of each of the 37 libraries tested. TPI1343, which has a pyrrolidine pentamine scaffold, showed the highest inhibition level and was selected for further studies. The general structure of the compounds in TPI1343 is shown in Fig. 3. It is composed of 120 mixtures, each containing one of the four functionalities defined and totaling 738,192 compounds and was synthesized using the synthesis scheme shown in Scheme 1 (Materials and Methods and Fig. S1, supplemental material) [29]. The 120 different systematically formatted mixture samples were synthesized using 26 different amino acids (R1,R2 and R3) and 42 different carboxylic acids (R4) in order to incorporate the R group functionalities. After the synthesis was completed, 120 different mixture samples of pyrrolidine pentamine analogs were obtained and screened (Fig. 3).

**Fig. 1.**
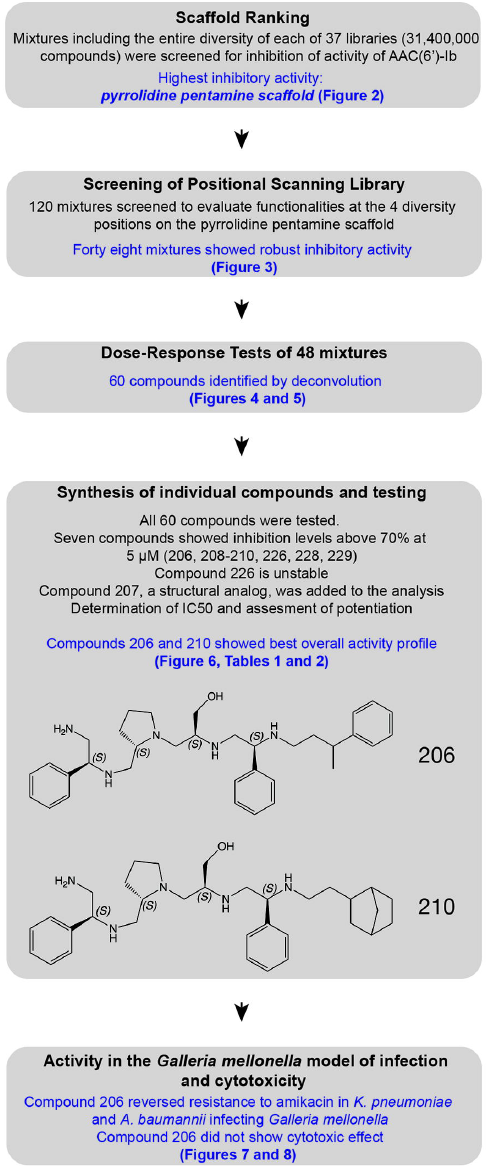
Screening strategy for identification of inhibitors of the AAC(6’)-Ib enzyme from combinatorial libraries. Thirty-seven libraries were tested to select the most potent scaffold, pyrrolidine pentamine. Next 120 mixtures were screened to identify functionalities at the 4 diversity positions. Forty-eight mixtures showed robust inhibitory activity. Deconvolution identified 60 individual compounds, which were synthesized in parallel. After preliminary characterization, two candidates, **206** and **210**, were singled for purification and further characterization.

**Fig. 2.**
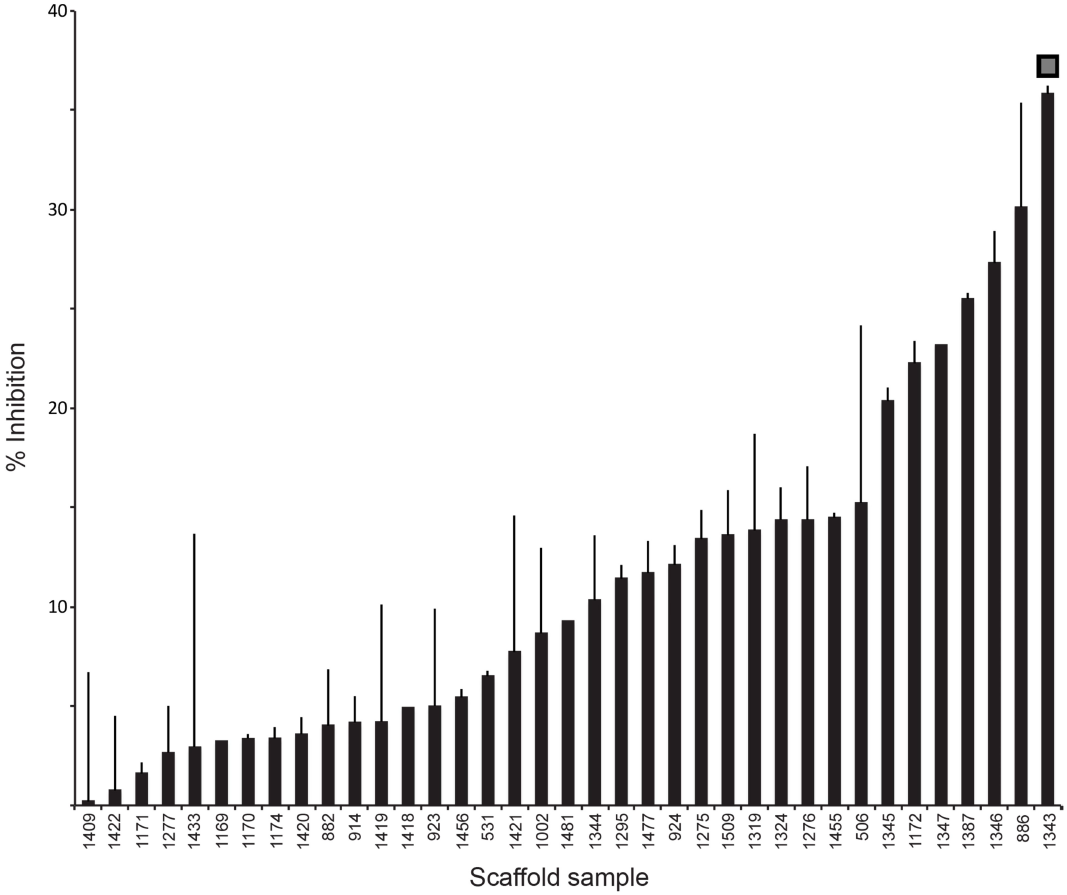
Scaffold ranking. Each mixture was added to the enzymatic reaction solution at 25 µg/ml. The activity of the enzyme with addition of the DMF was considered 100% and the percentage of inhibition was calculated based on activities in the presence of each mixture. Each assay was carried out in triplicate. The scaffolds corresponding to each TPI library are: 1409, Triazinetrione; 1422, N-Methyltriamine; 1171, Bis-cycle thiourea; 1277, Guanidino hydantoin; 1433, Nitrosamine; 1169, Bis-cyclic guanidine; 1170, Bis-diketopiperazine; 1174, N-acylated Bis-piperazine; 1420, N-methylated 1,3,4-trisubstituted piperazine; 882, C-6-acyloamino bicyclic guanidine; 914, N-Acyl triamines; 1419, N-Benzyl-1,4,5-trisubstited-2,3-diketopiperazine; 1418, N-Methyl-1,4,5-trisubstituted-2,3-diketopiperazine; 923, H-Tetrapeptide-NH2; 1456, Tetraamine; 531, Bicyclic Guanidine; 1421, N-benzylated 1,3,4-trisubstituted piperazine; 1002, Urea-linked bicyclic guanidine; 1481, Poly-phenylurea; 1344, Pyrrolidine Bis-diketopiperazine; 1295, Acylated cyclic guanidine; 1477, Platinum tetraamine; 924, Ac-Tetrapeptide-NH2; 1275, Dihydroimidazolyl-butyl-diketopiperazine; 1509, 2-imino-1,3,5-triazino [1,2-a] benzimidazoles; 1319, Dihydroimidazolyl-butyl-cyclic urea; 1324, Dihydroimidazolyl-methyl-diketopiperazine; 1276, Dihydroimidazolyl-butyl-cyclic thiourea; 1455, H-Tripeptide-NH2; 506, Reduced Dipeptidomimetics; 1345, Pyrrolidine Bis-piperazine; 1172, Bis-piperazine; 1347, Pyrrolidine Bis-cyclic thiourea; 1387, Trisubstituted triazinobenzimidazolediones; 1346, Pyrrolidine Bis-cyclic guanidine; 886; Benzothiazepine; 1343, Pyrrolidine pentamine.

**Fig. 3.**
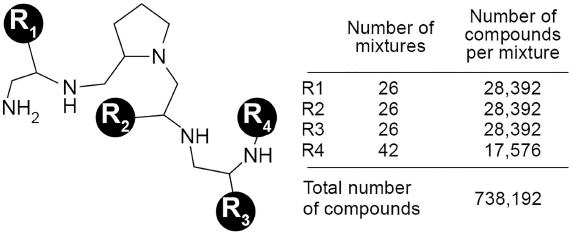
Pyrrolidine pentamine scaffold library TPI1343. The variable functionalities R1 – R4 are circled. The number of functionalities for each R group, the number of compounds per mixture, and the total number of compounds are shown to the right.

Forty-eight mixtures showed 70% or higher inhibitory activity (gray boxes, Fig. 4) and were selected for further dose-response tests at concentrations of 50, 25, and 12.5 µg/ml. Based on the results of these tests, the functionalities of the most active mixtures were selected for deconvolution, which resulted in 60 individual compounds (indicated by a bar in Fig. 5 and listed in Table S2, supplemental material). All 60 compounds were then synthesized and tested individually at concentrations 5 and 1 µM to determine their inhibitory activity of the AAC(6′)-Ib enzyme. Seven compounds showed the inhibition levels above 70% at 5 µM (**206**, **208**-**210**, **226**, **228**, **229**; see Table 1 and Table S2, supplemental material). One of them, compound **226**, could not be further analyzed because of possible chemical instability. The remainder were purified along with the close structural analog **207,** and used to determine their inhibitory characteristics. Table 1 shows the half-maximal inhibitory concentration (IC_50_) values and structures of these compounds. IC_50_ values rank between 2.1 µM (compound **206**) and 12.4 µM (compound **228**). Kinetic analysis to determine the mode of inhibition of all 7 compounds showed uncompetitive inhibition with respect to acetyl CoA and mixed inhibition with respect to kanamycin A (Fig. 6 and Fig. S2, supplemental material).

**Fig. 4.**
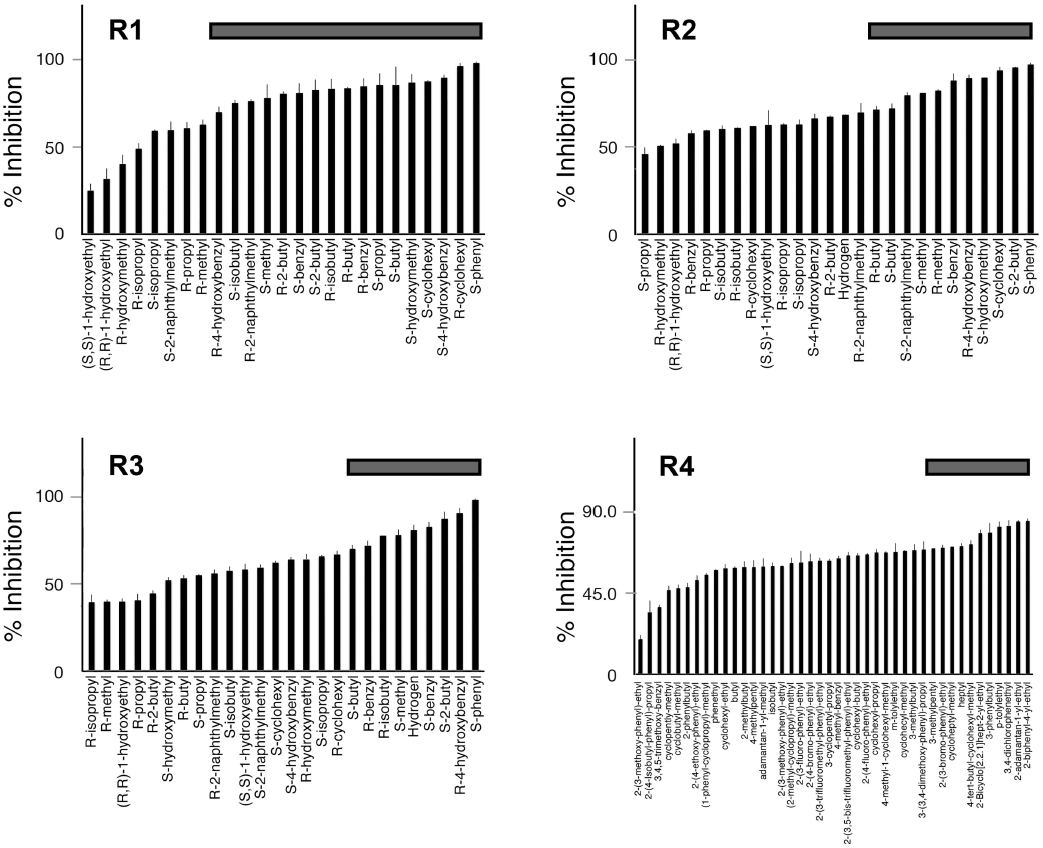
Positional scanning library screening results, ranking and selection. Each mixture was added to the enzymatic reaction solution at 50 µg/ml. The activity of the enzyme with addition of the DMF was considered 100% and the percentage of inhibition was calculated based on activities in the presence of each mixture. The defined functionalities in each mixture for each R group are indicated. The gray bars on top indicate the mixtures selected for dose-response assays.

**Fig. 5.**
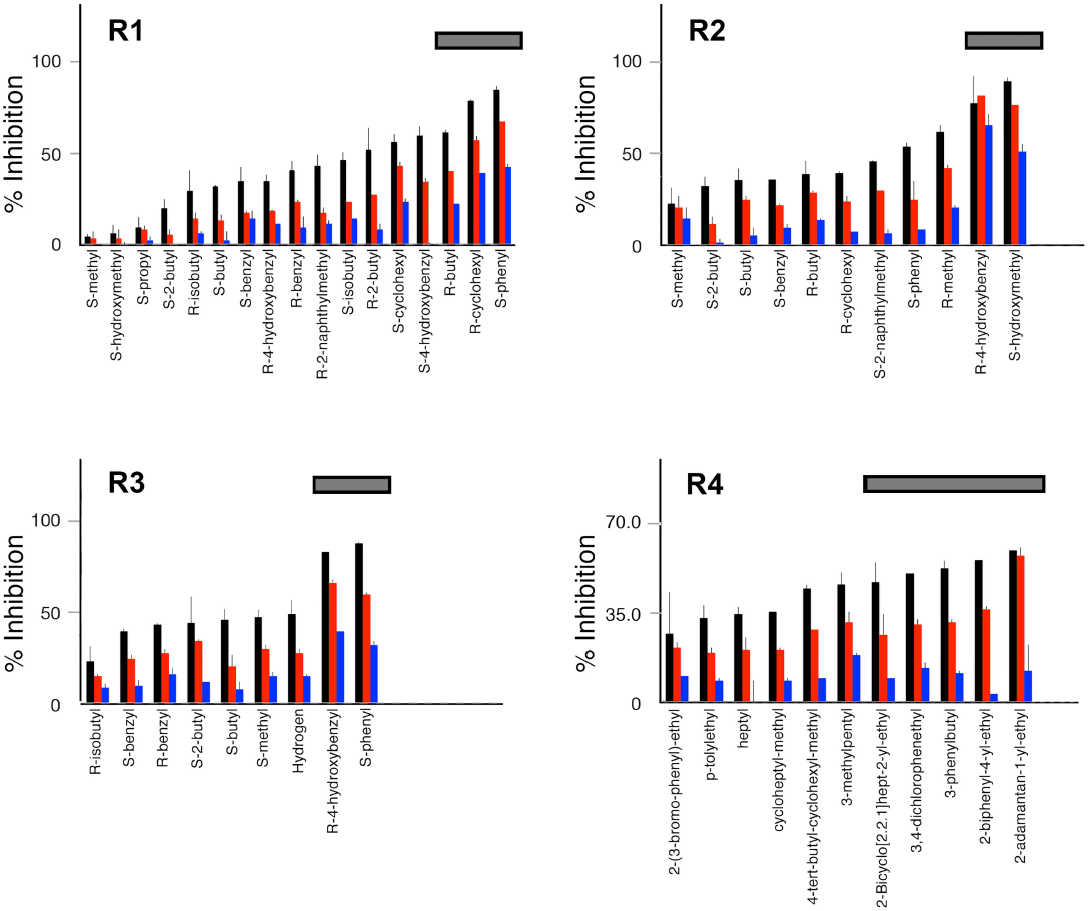
Dose-response assays. The mixtures selected on the basis of the results shown in Fig. 3 were added at 12.5 µg/ml (blue), 25 µg/ml (red), and 50 µg/ml (black) to the enzymatic reactions. The inhibition levels were calculated as in the previous assays. The gray bars at the top indicate the mixtures used for deconvolution of single compounds to be tested.

**Fig. 6.**
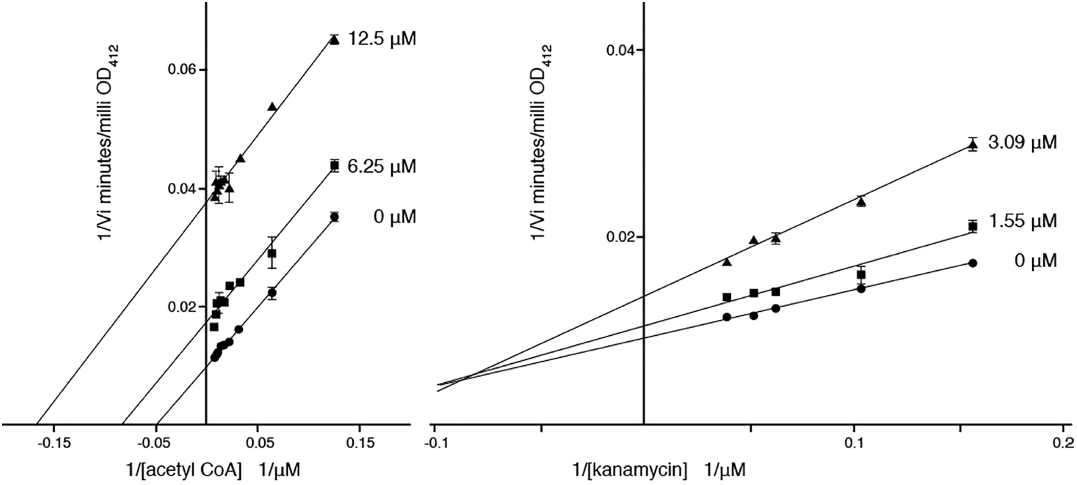
Inhibition of AAC(6’)-Ib by compound **206**. Lineweaver–Burk plots obtained carrying out reactions containing variable concentrations of acetyl CoA (left) and kanamycin A (right).

The compounds showing the lowest IC_50_ values (compounds **206**-**210**) are structural analogs that differ only in the R4 position. To assess these compounds’ ability to potentiate the activity of amikacin, checkerboard assays were performed with these five compounds with *E. coli* TOP10(pNW1), *A. baumannii* A155 and *K. pneumoniae* JHCK1. Compounds **207** and **208** showed the highest antimicrobial activity in the absence of amikacin in the *E. coli* (with IC_50_s of 12 and 16 µM, respectively) and *A. baum*annii (with IC_50_s of 12 µM each) strains, and did not show consistent potentiation in any of three bacteria tested. Compound **209** also showed antimicrobial activity in the absence of amikacin (with an IC_50_ of 20 µM in A155 and 8 µM in *E. coli*) but also showed 10-fold potentiation of the amikacin IC_50_ in *A. baumannii* A155 and 3-fold potentiation of the amikacin IC_50_ in *K. pneumoniae* JHCK1. Compounds **206** and **210** showed the best activity profile of all, with IC_50_ values greater than 24 µM in all strains and potentiation of 3-to 6-fold of the amikacin IC_50_ in all strains (Table 2 and Table S3, supplemental material).

**Table 2.**
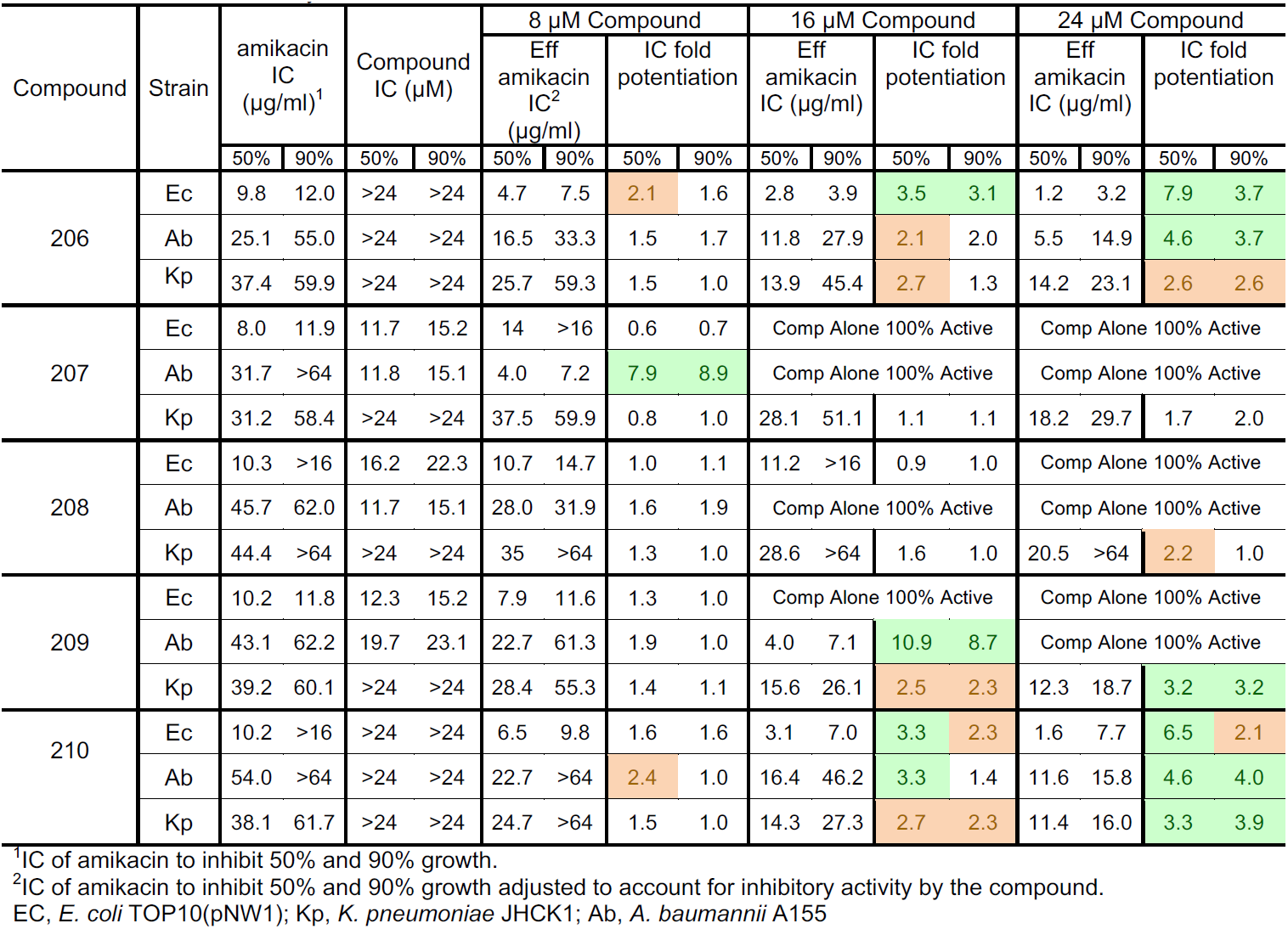
Checkerboard assays

The three most favorable compounds from the checkerboard assay (**206**, **209**, and **210)** were then further assessed for their potential capability to be used as therapeutic agents to treat amikacin-resistant infections using the *G. mellonella* virulence model. This model of infection has been extensively validated for the study of host-pathogen interactions as well as to assess the efficacy of new antibiotic treatments against several bacterial pathogens [34]. Infection with *A. baumannii* A155 or *K. pneumoniae* JHCK1 bacteria resulted in a significant increase in mortality compared with larvae that were not injected with the bacteria or were injected with sterile PBS, amikacin, or any of the compounds (Fig. 7A). No significant reduction in mortality was observed when the individuals were injected with bacteria plus amikacin or amikacin and compound **209** or **210** (Fig. 7B and C; LogRank test *p*>0.05 in all cases). Conversely, the group treated with amikacin plus compound **206** experienced a significantly reduced mortality rate (Fig. 7B and C; LogRank test *p*<0.05 for both strains). These results indicate that compound **206** is a good lead for further optimization to identify a potent inhibitor of the resistance to amikacin mediated by AAC(6’)-Ib.

**Fig. 7.**
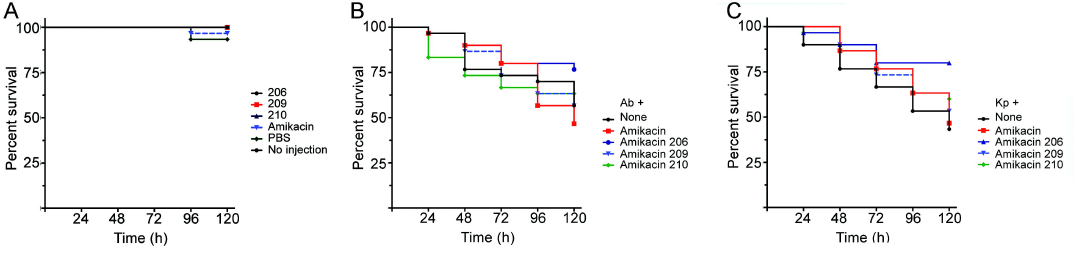
*G. mellonella* infection and treatment assays. Larva groups (10 individuals) were injected with the components shown in the figure. Panel A, larva were injected with the different components but they were not infected. Panels B and C, larva were infected with *A. baumannii* A155 (Ab, panel B) or *K. pneumo*niae JHCK1 (Kp, panel C), and the indicated components. The amikacin concentrations were 16 and 6 mg AMK/kg of body weight for *K. pneumo*niae JHCK1 and *A. baumannii* A155, respectively. The concentrations of compounds **206**, **209**, or **210** were 10 µM. The larvae were incubated at 37°C in the dark, and survival was recorded at 24-h intervals over 120 h.

*G. mellonella* has been used before as model for evaluating toxicity [35]. The infection assays using the *G. mellonella* model did not indicate that compound **206** was acutely toxic to the host at the concentrations used. To confirm this result, compound **206** was tested using a standard cytotoxicity assay as described in the Materials and Methods section. Addition of compound **206** to the cells at concentrations higher than those needed to observe its inhibitory activity did not result in significant toxicity (Fig. 8).

**Fig. 8.**
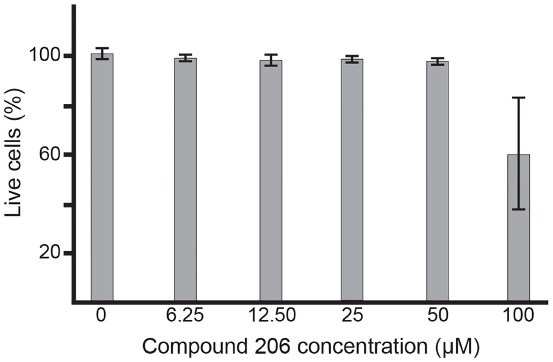
Cytotoxicity of compound **206**. Cytotoxicity towards HEK293 cells was tested using a LIVE/DEAD assay kit at the indicated concentrations of compound **206**. The percentage of dead cells was calculated relative to the cells treated with DMSO. Cells incubated with 0.1% Triton X-100 for 10 min was used as a control for maximum toxicity. Experiments were conducted in triplicate and the values are Mean ± SD.

## 4. Discussion

There are many mechanisms by which bacteria resist the action of aminoglycosides but the most significant is that of enzymatic modification mediated by aminoglycoside modifying enzymes [5]. Although there are more than hundred known enzymes, AAC(6’)-Ib is by far the most common cause of amikacin resistance [7].

A well-known approach to overcome resistance to antibiotics is the design of drugs that interfere with the resistance and when administered in combination with the appropriate antibiotic facilitate the treatment of otherwise resistant infections. This approach has been highly successful in the case of β-lactams, and there are currently several formulations composed of the antibiotic and the β-lactamase inhibitor in the market [36, 37]. In the case of aminoglycoside antibiotics, there were numerous reports identifying inhibitors of the expression of the aminoglycoside modifying enzyme as well as traditional enzyme inhibitors and compounds that may inhibit the enzymatic inactivation by protecting the antibiotic substrate molecule [37]. However, none of them could yet be developed for its use in the clinics.

To identify small molecule compounds that are inhibitors of AAC(6’)-Ib, we used mixture-based combinatorial libraries, the scaffold ranking approach, and the positional scanning strategy. Mixture-based small molecule libraries permit the testing of thousands of compounds by screening only hundreds of mixtures each of the scaffolds [24]. Potent inhibitors of biological processes have been identified following this approach. Examples are hexapeptides that inhibit recombination at the Holliday junction stage in the processes of integration or excision of the bacteriophage λ that permitted a better analysis of the recombination processes [38] and tetrapeptides that restored fully functional signaling and efficacy of polymorphic receptors that could result in a reduction of morbid obesity [39]. The scaffold ranking of mixture libraries indicated that there are several scaffolds that may include compounds with inhibitory activity towards AAC(6’)-Ib. The analysis of the scaffold ranking sample that was apparently the most potent, the pyrrolidine pentamine scaffold library consisting of 738,192 compounds with four diversity positions (R1-R4), led to the identification of compound **206** (R1=S-phenyl, R2=S-hydroxymethyl, R3=S-phenyl, R4=3-phenylbutyl).

AAC(6’)-Ib acts through an ordered kinetic mechanism where acetylCoA is the first substrate to bind [40]. Therefore, the uncompetitive inhibition exhibited by compound **206** indicates that it must bind the enzyme after acetylCoA. The mixed nature of the inhibition with respect to the aminoglycoside substrate suggest that compound **206** interferes with binding of the aminoglycoside and reduces the catalytic activity.

The screening of the scaffold ranking library (Fig. 2), the pyrrolidine pentamine positional scanning library (Figs. 4 and 5) and subsequent individual compounds derived from the library provide some structure activity relationship (SAR) data that can be utilized in future optimization studies. The scaffold ranking library screening identified the pyrrolidine pentamine scaffold as the most active inhibitor of AAC(6’)-Ib enzymatic activity. Screening of the positional scanning library produced good differentiation in inhibitory activity for the samples screened; this is an indication that specific individual compounds with key functionalities at each diversity position are driving the activity. From the positional scanning data the key functionalities appear to be a mixture of polar (eg., hydroxybenzyl) and non-polar (eg., phenyl) depending on the diversity position. The screening of the individual compounds confirms the importance of placing the right functional group in the correct diversity position as the selected individual compounds displayed a range of potencies (Figs. 4 and 5). Closer inspection of the individual compounds through sorting of the structural analogs (Table S4, supplemental material) reveals that the R1 (*S*-phenyl), R2 (*S*-hydroxymethyl), and R3 (*S*-phenyl) is a critical motif for activity within this series of compounds (**206** through **210**). R2 and R3 substitution analogs (compounds **201**, **205**, **216**, and **220**) displayed significantly reduced activity as compared to compounds **206** and **210**. The reduction in activity was less significant with the R1 analogs (compounds **226**, **230**, **246**, and **250**). Thus for AAC(6’)-Ib activity the R1 (*S*-phenyl), R2 (*S*-hydroxymethyl), and R3 (*S*-phenyl) has been identified as the critical structural motif, with substitutions made at the R4 position being more permissive. However, the antimicrobial activity in the absence and presence of AMK differed between these compounds. Compounds **206** and **210** did not have substantial antimicrobial activity alone, with MIC 50% and 90% >24 µM for *E. coli* TOP10(pNW1), *A. baumannii* A155 and *K. pneumoniae* JHCK1, in contrast, compounds **207**, **208** and **209**. Furthermore, compounds **206**, **209** and **210** were the compounds that potentiated AMK antimicrobial activity. In addition, compound **206** in combination with AMK was the only compound that showed reduced mortality in the *G. mellonella* virulence model treated with amikacin-resistant infections. In future studies, we will explore further the importance and effect of R4 substitutions for enzyme activity as well as potentiation in checkerboard assay. These studies could provide additional SAR data on AAC(6’)-Ib inhibitory activity and potentiation and will be part of the medicinal chemistry efforts to produce a preclinical candidate for AAC(6’)-Ib to treat drug resistant bacterial infections. Furthermore, identification of these and other inhibitors could provide tools for the analysis and better understanding of the AAC(6’)-Ib enzymatic mechanisms.

## 5. Conclusions

Using mixture-based combinatorial libraries, the scaffold ranking approach, and the positional scanning strategy we identified an inhibitor of the AAC(6’)-Ib enzyme. Inhibitors of resistance enzymes could be used for analysis of the properties of the enzymes and molecular mechanisms of the reaction and, after further development and enhancement of activity for clinical use to overcome resistance.

## Declarations

### Funding

This work was supported by Public Health Service grant 2R15AI047115-04 (MET) from the National Institute of Allergy and Infectious Diseases, National Institutes of Health and the State of Florida, Executive Office of the Governor’s Office of Tourism, Trade, and Economic Development.

### Competing Interests

None

